# Learning predictive signatures of HLA type from T-cell repertoires

**DOI:** 10.1101/2024.01.25.577228

**Authors:** Maria Ruiz Ortega, Mikhail V. Pogorelyy, Anastasia A. Minervina, Paul G. Thomas, Aleksandra M. Walczak, Thierry Mora

## Abstract

T cells recognize a wide range of pathogens using surface receptors that interact directly with peptides presented on major histocompatibility complexes (MHC) encoded by the HLA loci in humans. Understanding the association between T cell receptors (TCR) and HLA alleles is an important step towards predicting TCR-antigen specificity from sequences. Here we analyze the TCR alpha and beta repertoires of large cohorts of HLA-typed donors to systematically infer such associations, by looking for overrepresentation of TCRs in individuals with a common allele.TCRs, associated with a specific HLA allele, exhibit sequence similarities that suggest prior antigen exposure. Immune repertoire sequencing has produced large numbers of datasets, however the HLA type of the corresponding donors is rarely available. Using our TCR-HLA associations, we trained a computational model to predict the HLA type of individuals from their TCR repertoire alone. We propose an iterative procedure to refine this model by using data from large cohorts of untyped individuals, by recursively typing them using the model itself. The resulting model shows good predictive performance, even for relatively rare HLA alleles.

## I. INTRODUCTION

T cells are one of the building blocks of adaptive immunity. They accomplish a highly precise and tailored immune response using membrane-bound antigen-specific T cell receptors (TCR), a heterodimer composed of an alpha and a beta chain. To cover all possible pathogens, T cells rely on their large TCR sequence diversity [1]: the gene of each chain undergoes somatic V(D)J recombination during T cell development, resulting in the generation of extensive genetic heterogeneity in the TCR repertoire.

The *αβ* TCR interacts with short peptide fragments displayed on the surface of antigen presenting cells by the major histocompatibility complex (MHC) molecule. In humans, these structures are encoded by the highly polymorphic human leukocyte antigen (HLA) genes. Each individual carries up to 2 alleles of each of the 3 class-I (A, B, and C) and multiple class-II (DP, DQ, and DR) MHC genes, each binding a large but restricted set of different peptides with common sequence features [2–4]. This MHC restriction introduces an extra layer of complexity to TCR specificity to particular combinations of peptides and MHC molecules [5, 6], with strong heterogeneity across individuals. Despite its importance for antigen specificity, the interaction between HLA type and the TCR repertoire remains poorly understood, especially for less common HLA alleles.

Advances in high-throughput repertoire sequencing [7] have allowed for unprecedented insights into the composition of immune repertoires, which constitutes a dynamic register of the immune challenges encountered by the organism [8]. Public TCR clones, which are found in several individuals, have raised great interest as possibly playing functional roles in antigen recognition [9, 10]. Many factors affect TCR sharing, including biases in the VDJ recombination process and convergent recombination [11–15], thymic and peripheral selection [16], as well as shared diseases and HLA background [8, 17]. Here we analyze the TCR repertoire of 1039 HLA-typed human donors from three different cohorts. Following [8, 17] we identify public TCR sequences that are enriched in individuals with a given HLA allele. We use those sequences to build a reliable predictor of HLA type from repertoire data. Our analysis combines alpha and beta chain information, and makes predictions for both class-I and II HLA alleles, leading to more accurate and comprehensive predictions than previous approaches. We demonstrate how our HLA type predictor, encoded in new software package, HLAGuessr (available at https://github.com/statbiophys/HLAGuessr), can be used to discover new HLA-TCR associations from the repertoires of untyped individuals only, opening up the possibility to aggregate data from a large body of datasets for which the conventional HLA typing is unavailable.

## II. RESULTS

### Disovery of HLA-associated TCRs

We first sought to identify lists of TCRs associated with each HLA allele, using an aggregate dataset of TCR repertoires of 1039 HLA-typed donors from three different sources comprising: the *Russell et al*. dataset with 718, 000 unique TCR*α* and 1, 016, 000 unique TCR*β* sequences from 237 donors [18], the *Emerson et al*. dataset with more than 7 *×* 10^7^ TCR *β* unique sequences from 610 donors [17] and the *Rosati et al*. dataset with 7 *×* 10^6^ TCR*α* and 1 *×* 10^7^ TCR*β* sequences from 192 healthy and Crohn’s disease patients [19]. We only considered TCR sequences that were seen in at least 3 people across the cohorts, leaving us with 654,535 alpha chains, and 4,927,204 beta chains to analyse. For each individual, the HLA type was encoded in a list of up to 6 HLA type I alleles (2 of each of A, B, C) and at least 10 for HLA type II (4 possible isoforms of DP, 4 of DQ and 2 of DR1, while DR3-5 have variable presence in humans). There were 316 HLA alleles in total.

For each TCR sequence (alpha or beta), and each HLA allele, we performed Fisher’s two-tailed exact test to ask whether the presence of that TCR in individuals was associated with the presence of that allele (see Methods for details) in each cohort. To control for multiple testing, we set the p-value threshold for a significant association using the Benjamini-Hochberg procedure, so that the false positive rate is 5%. This procedure yields a list of TCR*α* and TCR*β* sequences that are associated with a particular HLA allele, with a level of significance indicated by the p-value (see Supplementary Material). The results for each of the three cohorts analysed separately are summarized in the Venn diagrams of Fig. 1A, where the overlap refers to TCR clonotypes that were associated with the same HLA in different cohorts. We found associations in the *Russell et al*. cohort between 894 TCR*α* with 51 HLA alleles, and between 863 TCR*β* and 58 HLA alleles. In the *Emerson et al*. cohort, we report 17889 TCR*β* associated to 116 HLA alleles; this number broadly agree with the previous analysis of [8] with minor discrepancies due to a different definition of alleles: we restrict our analysis to single HLA alleles, while [8] also considered haplotypes or co-occurence of several alleles. In the *Rosati et al*. cohort, we find 384 TCR*α* associated with 48 HLA alleles, and 1147 TCR*β* associated to 58 HLA alleles. The number of significant associations is largely determined by the number of sequences in each dataset, which explains large differences between the cohorts.

**FIG. 1.**
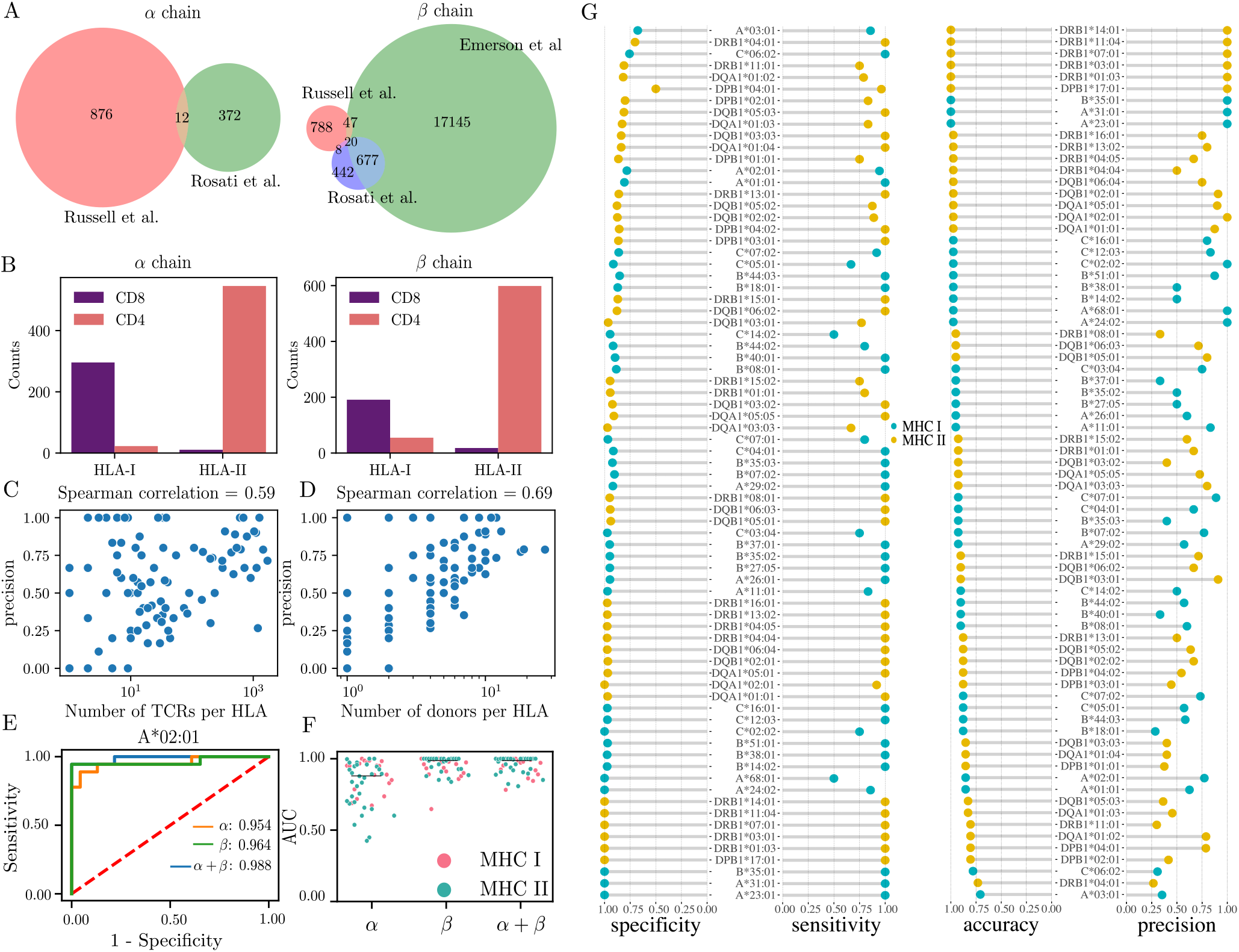
Summary of HLA predictor performance. (A) Venn diagram of the alpha and beta chains significantly associated with an HLA allele after performing the Fisher’s exact test in our three different cohorts of data. Overlap corresponds to sequences associated to the same HLA in the different datasets. (B) The number of HLA-associated TCRs show concordance between their CD4+ and CD8+ phenotype and the MHC class of the HLA allele. (C-D) Correlation between the precision of the classifier with (C) the number of found associations and (D) the frequency of the HLA allele in the cohort. (E) Receiving Operator Characteristic (ROC) of the classifier for the A*02:01 allele in the validation data set, using either each chain separately or together. (F) Area under the curve (AUC) of the ROC for each of the tested alleles. The average value of each group is indicated with the horizontal black line. (G) Performance metrics of the HLA classifier on the validation dataset. Specificity=TN/(TN+FP), Sensitivity=TP/(TP+FN), Accuracy=TP+TN/(TP+TN+FP+FN), Precision=TP/(TP+FP), where T/F stand for true/false, and P/N for positives/negatives.

As a first check of the consistency of the discovered associations, we verified that the CD4/CD8 phenotype of each TCR reported in the *Russell et al*. dataset (as determined by flow cytometry) matched their associated HLA class II and class I alleles. CD4^+^ cells recognize antigens presented on class II MHC molecules (encoded by the DP, DQ, and DR HLA genes), while CD8^+^ respond to antigens presented on class I MHC molecules (encoded in the A, B, and C HLA genes) [20]. Fig. 1B shows the number of CD4 and CD8 TCR *α* and *β* sequences that were associated to any HLA of class I or II. As expected, the majority of CD8^+^ TCRs are associated to HLA alleles of class I, and the majority of CD4^+^ TCRs to HLA alleles of class II. Exceptions to this rule are not necessarily all due to noise, since some sequences are simultaneously marked as CD4 and CD8 cells, probably because the same alpha or beta chain may be used in two distinct T cell clones with different HLA specificities.

### HLA-associated TCR predict the HLA type

To maximize statistical power, we repeated the association discovery procedure on a random 70% training portion of the merged dataset combining all three cohorts, resulting in a list of 2570 TCR*α* and 63060 TCR*β* sequences significantly associated to at least one HLA allele. This number is much larger than the sum of the numbers of associations from each cohort taken separately. For each HLA allele, we fit a logistic regression model on the training set predicting whether a given donor has this allele or not, depending on whether it has or not each TCR sequence (both alpha and beta) found to be associated to that allele. We excluded from this fit sequences from the *Emerson et al*. dataset whenever the alpha chains were used. Regression was done with an L2 regularization, which strength was tuned to optimize the accuracy of the prediction on the testing set made of the remaining 30% portion of the data (see details in Methods).

Model predictions were then evaluated on an independent dataset of 46 HLA-typed individuals from *Milighetti et al* [21]. Four measures of performance for each analysed HLA allele are reported in Fig. 1G: accuracy (proportion of individuals for which the prediction is correct), precision (proportion of true positives among positive predictions), specificity (propotion of people without the HLA allele who are correctly predicted), and sensitivity (proportion of people with the HLA allele who are correctly predicted). The accuracy and specificity are good for almost all alleles, due to the fact that the prevalance of each HLA allele is relatively low in the population, meaning that there many more negative than positive examples. We expect that HLA alleles with lower prevalence are harder to predict [22–24]. Indeed, precision is correlated with both the total number of TCR associated to the HLA allele (Pearson *ρ* = 0.59, *p* = 5 *×* 10^*−*17^, Fig. 1C), and the prevalence of that HLA allele in the cohort (Pearson *ρ* = 0.69, *p* = 5 *×* 10^*−*12^, Fig. 1D). A more complete way to report the performance of the classifier is to plot the Receiver Operating Characteristic (ROC) curve, which shows sensitivity as a function of specificity as one varies the threshold on the logistic regression score (this threshold was set earlier by the logistic regression to maximize cross-entropy, see Methods). Example ROC curves are shown in Fig. 1E for HLA A*02:01, using either both alpha and beta chains, or each separately. The ROC curve may be conveniently summarized in a threshold-independent manner by the area under the curve (AUC), which is 1 if the predictor is perfect, and 0.5 if it is random. Fig. 1F shows the AUC of all HLA alleles, comparing predictors using each chain separately, or both of them together. Although TCR*α* and TCR*β* could be used for HLA typing independently, the beta chain (AUC=0.96 *±* 0.06) is more informative than the alpha chain (AUC= 0.84 *±* 0.15, *p* = 6 *×* 10^*−*8^, Wilcoxon test). Using both chains together does not confer a significant advantage compared to the beta chain alone (AUC=0.94 *±* 0.11, *p* = 0.3).

### Fuzzy TCR sequence matching improves HLA-typing

We also tested two variants of the algorithm. Since TCR specific to the same target often have similar sequences [17, 25–28], one could extend the search for the presence of HLA-associated TCRs to their neighbors in sequence space. We re-trained the logistic regressor by counting each HLA-associated sequence as present if either that exact sequence or a variant with one amino acid mismatch in CDR3 was found in the repertoire. Fig. 2A shows that this more liberal way to include HLA-associated sequences leads to increased performance on average, as measured by the AUC (0.95 *±* 0.05, *p* = 3 *×* 10^*−*4^ Wilcoxon test). In the second variant, which is equivalent to the original method of Ref. [17] and requires no logistic regression, the prediction of whether the individual is positive or negative for an HLA is just determined by the total number of HLA-associated sequences found in their repertoire. Fig. 2B shows this simpler strategy typically worsens the quality of the prediction (AUC= 0.90 *±* 0.12, *p* = 2 *×* 10^*−*5^ Wilcoxon test), emphasizing the importance of logistic regression in which the contributions of distinct TCRs to the prediction score are different (see (Fig. S1)). Thus both inexact matching and weighting individual TCR sequence improve HLA prediction quality, but the former is more computationally expensive. For this reason, we used the basic classification algorithm in further analysis.

**FIG. 2.**
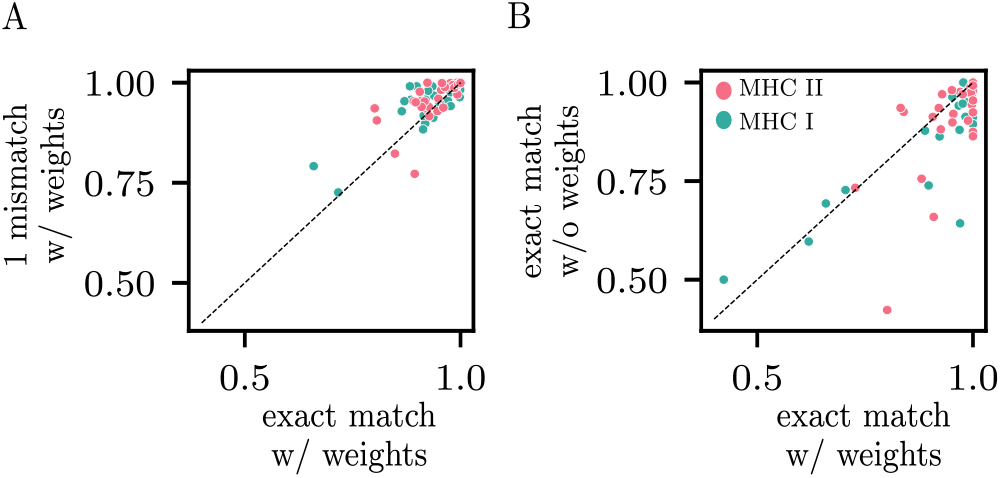
Algorithm variants. (A) Comparison of AUCs between the basic classifier (based on exact matches with HLA-associated TCR), and a classifier where single amino acid variants are counted as a match. (B) Comparison of AUCs between the basic classifier, where the weights associated to each HLA-associated TCR are learned using logistic regression, and a simpler classifier that simply counts the number of matches (without weights, i.e. all weights=1), which was used in [17].

### Features of HLA-associated TCRs

We then asked whether HLA-associated TCRs share some common features, focusing on HLA A*02:01, the most abundant HLA allele in our dataset. Fig. 3A shows the CDR3 amino acid length distribution for both alpha and beta chain (28 and 221 sequences, respectively). The CDR3-length distribution of the beta chain of HLA A*02:01-associated sequences seems to be heavily skewed towards shorter sequences. *α* chains strongly deviates as well when compared with random TCRs (Fig. 3A). Since TCR:pMHC binding takes place through the V gene-encoded CDR1 and CDR2 contacting the conserved helical residues of the MHC [29], we expect that some conserved structural footprint might be observed in V gene usage. A*02:01-associated TCRs showed a biased usage of V genes relative to generic TCRs, with enrichment of TRBV10, TRBV19, TRBV29, and under-representation of TRBV15,TRAV24,TRAV5 V gene families (Fig. 3B. We reasoned that the usage of those genes may be biased in the whole repertoire (not restricted to the HLA-associated TCR) of A*02:01 positive individuals as well.

**FIG. 3.**
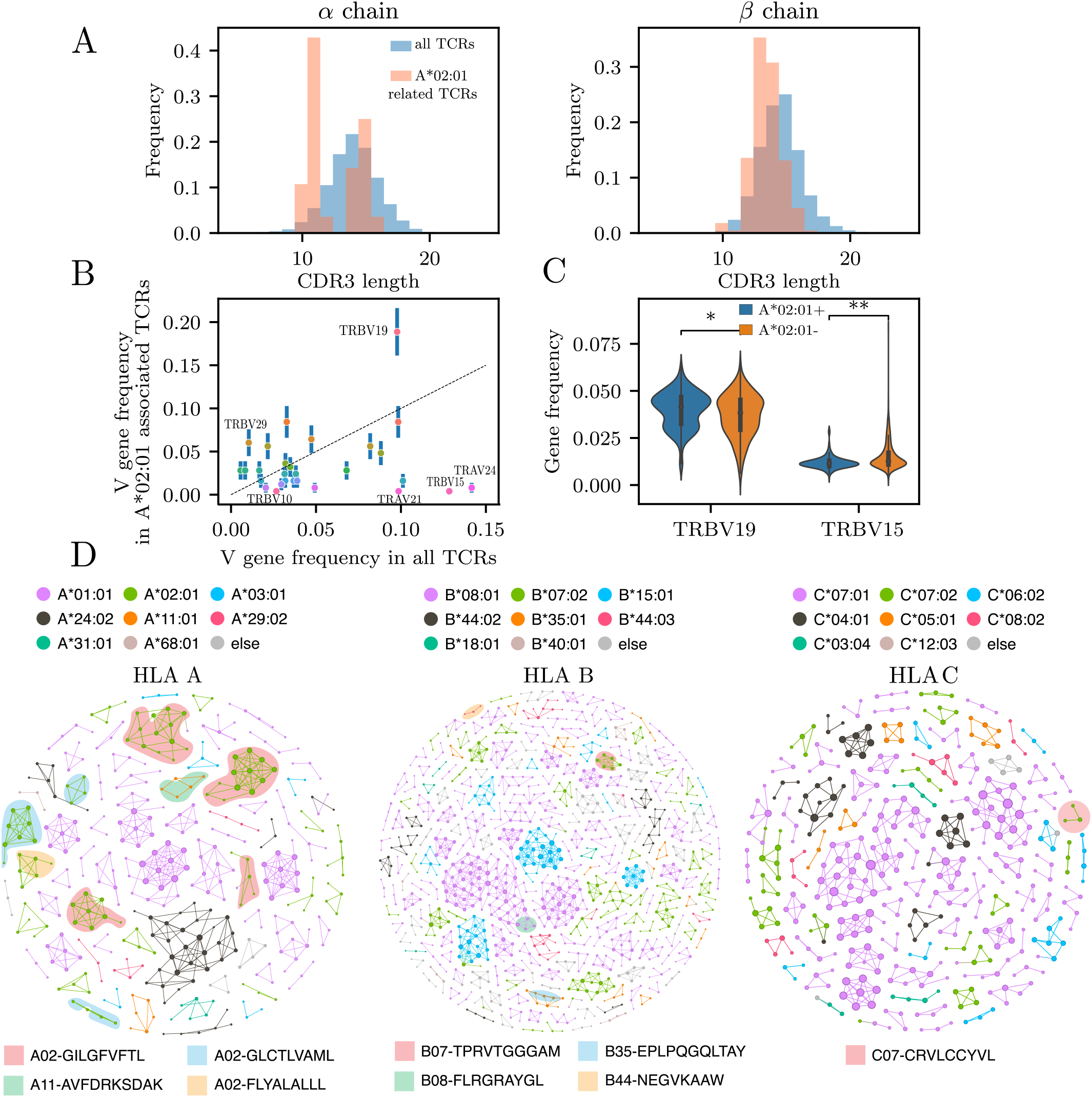
Features of the HLA-related sequences. (A) CDR3 length distribution of A*02:01-associated TCRs, compared to all TCRs. (B) V gene frequency usage comparison between the same two subsets. Some V gene families are preferentially used in A*02:01-related TCRs (TRBV10, TRBV19, TRBV29) while other genes are underrepresented (TRBV15,TRAV24,TRAV21). (C) Whole-repertoire frequencies of two representative differentially used genes, TRBV19 and TRBV15, in A*02:01 positive vs negative individuals show small but statistcally significant difference in those genes. (D) Network analysis of TCR*β* chains associated with different A, B and C HLA-alleles (TCR*α* were excluded since they formed negligible networks). Each node represent an amino acid CDR3 + V gene clonotype. Edges connect clonotypes that have at most on amino acid mismatch but the same length and V gene. Each color represent the specific HLA to which the TCR is responsive. Shadow color correspond to epitopes for which those TCRs are found to be responsive from the VDJdb database [30].

We found significant differences between the usage of TRBV19 and TRBV15 (the two genes with the largest deviation) of A*02:01 positive versus negative individuals (*p* = 1 *×* 10^*−*4^ and *p* = 6 *×* 10^*−*17^, Mann-Whitney U test, see Fig. 3C). This suggests that A*02:01-associated TCRs biases towards certain V genes could affect the repertoire of A*02:01 positive people as a whole.

Next, we investigated if similar TCRs are associated with the same HLA alleles, as already suggested by the improved performance of the mismatch-tolerant version of the HLA-type predictor (Fig. 2A), where similar TCR to the ones actually associated to the HLA also help determine if an individual is positive for that HLA.

In Fig. 3D, we represent HLA class I-associated TCRs as similarity networks, where nodes represent TCR clonotypes, defined by the CDR3 sequence and the V gene family, and edges are drawn between two sequences of the same length that differ by at most one amino acid. Each HLA allele may be represented by several clusters, corresponding to distinct specicifity classes within the same HLA, and some of these specificity classes can be mapped out onto specific antigens by matching clonotypes from the VDJdb database [30]. We downladed human TCRs from VDJdb (accessed on 22 September 2023), and excluded sequences from the 10x Genomics highly multiplexed dextramer dataset [31]. We then searched for exact matches of both CDR3 amino acid sequence and V gene family. Almost all clusters associate with a single HLA allele, with no mixing (Fig. 3D). This suggests that during thymic selection and further clonal expansion driven by antigen recognition each HLA selects exclusive clusters of closely related TCR sequences.

The number of memory T cells progressively increases with age due to antigen exposure [32, 33], and we expect the repertoire to focus on specific HLA types during that process. We asked how the number of HLA-related TCRs in each individual from *Britanova et al*. changes with age. We found that the relative number of HLA-related TCRs found in repertoires from individuals from 0 to more than 100 years old shows a weak yet significant correlation with age (*r* = 0.29, *p* = 0.01, Fig. S2).

### Predicting HLA alleles in untyped repertoires

Our HLA predictor was trained on the repertoires of individuals whose HLA type was known. However, there exist many more datasets for which this information is absent. We asked if we could leverage this large amount of untyped repertoire data to improve the performance of the classifier in an iterative way, by first typing untyped repertoires computationally using our logistic predictor, and then use these repertoires to discover new TCR-HLA associations. To demonstrate the feasibility of this approach, we used two additional TCR*β* repertoire datasets: one comprising 108 healthy individuals from Ref. [34], and the second including 1414 donors with a confirmed SARS-CoV-2 infection at various timepoints following the peak of the disease, totaling 1.6 *×* 10^8^ reads [35], and focused on 4 carefully selected HLA alleles. We picked A*02:01, DPB1*04:01 and DQB1*01:02 because of their high prevalence (around 50%) in the population, which allows for more robust statistics. The initial classifier showed only moderate accuracy for DPB1*04:01 and DQB1*01:02 despite their abundance, suggesting room for improvement. We chose the 4th allele DPB1*02:01 because of its mediocre performance in the initial classifier (AUC of 0.77), again to test the potential for improvement of an iterative approach.

We divided the cohort of untyped individuals into random subgroups of 100. We then applied the following recursion to each of the 4 HLA alleles. We first used the classifier previously trained on the typed cohort to predict the HLA positive or negative status of the first sub-group of 100. We then added this newly typed subgroup to the initial cohort, and re-ran the Fisher’s exact test to obtain a new list of HLA-associated TCR sequences. This updated list of HLA-associated sequences is expected to be longer than the previous one, as it is based on a larger cohort. We use this list to re-train a logistic regression classifier on the augmented cohort (initial cohort plus the 100 subgroup), using again a 70%/30% split between training and testing to optimize the regularization parameter. The new classifier is used to type the next subgroup of 100 individuals. The procedure is repeated until all subgroups have been typed.

With each iteration and the inclusion of more individuals, the statistical power is increased, leading to an exploding number of HLA-associated TCR, notably after the 10th iteration (see Fig. 4A). However, since there are errors in the computational typing, one might worry that these errors get amplified and propagated through the iterations. In order to mitigate this effect and also to avoid overfitting due to a very large number of associated sequences, we tune the threshold on the p-value of the Fisher’s exact test so that the number of HLA-associated sequences is proportional to the number of included donors, with a ratio fixed to its value found in the initial cohort.

**FIG. 4.**
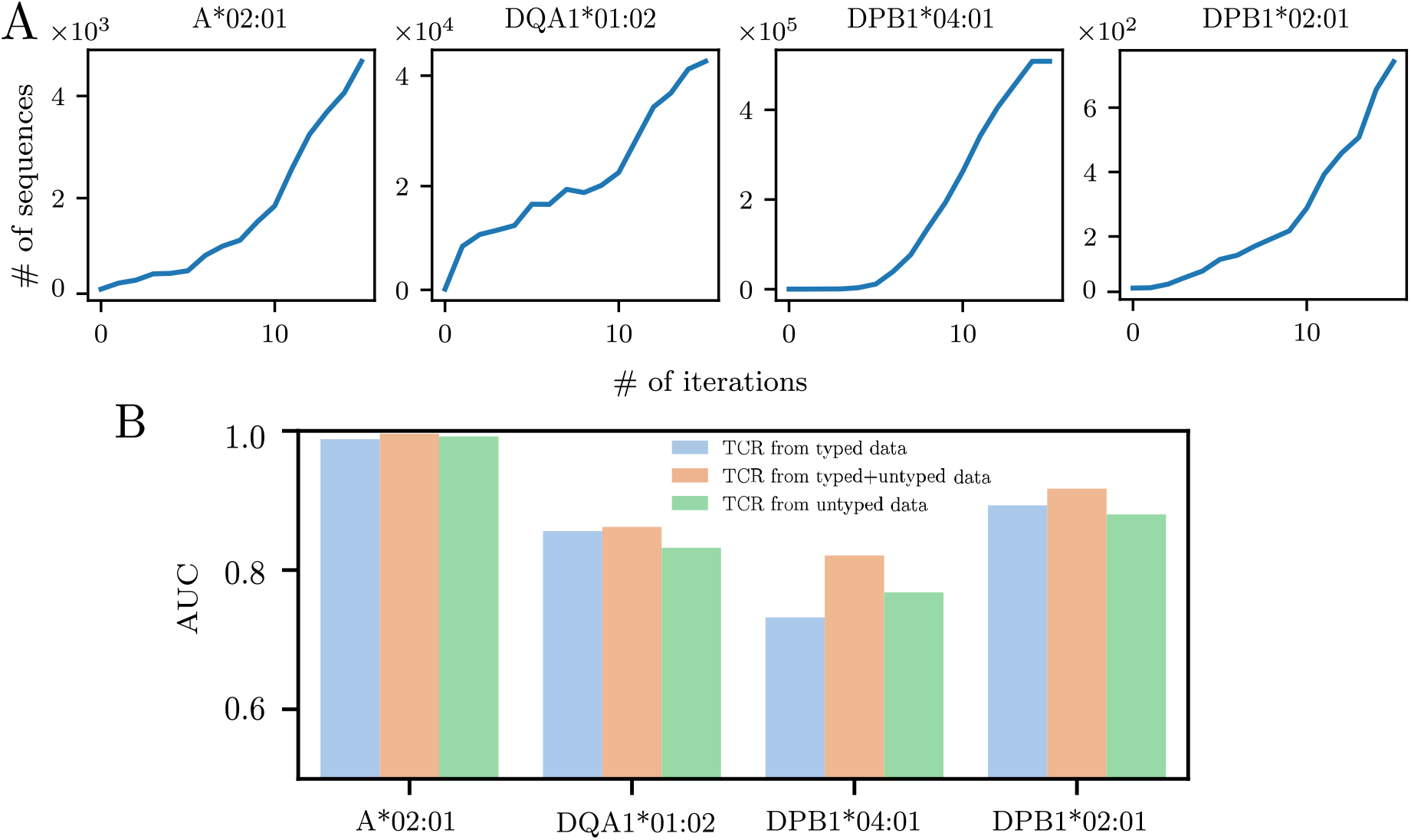
Iterative HLA-typing process. (a) Number TCRs found to be significantly related to the HLA alleles A*02:01, DQA1*01:02, DPB1*04:01, DPB1*02:01 after recursively applying our HLA predictor in two untyped datasets [34, 35]. In this procedure, donors are iteratively typed by groups of 100 patients. The HLA related sequences found in our newly typed subcohort are used to infer the HLA type of the next 100 patients. (b) Comparison of the AUC when predicting in the validation dataset using the sequences inferred in the first round from our initial typed cohort (blue bar) the ones learned exclusively from the untyped cohorts after 14 iterations (green bar) or the combination of both subcohorts (orange bar). The performance is shown to be maximized when using all the available HLA-related TCRs, supporting the quality of the information acquired through this iterative process.

Fig. 4B shows that the performance (AUC) of the classifier on the validation cohort is improved in the iterative procedure (orange) compared to the original classifier (blue), suggesting that the new associations provide additional information. To further validate the new associations, we trained a third logistic classifier using the presence or absence of the novel HLA-associated TCR only, i.e. sequences that were not present in the original list. This classifier yielded comparable and sometimes even better performance (in green) than the original classifier, despite being based on sequences that were identified using untyped cohorts. This demonstrates the potential of the iterative method to uncover novel associations with strong predictive power.

## III. DISCUSSION

Despite the advances in the high-throughput sequencing technology of immune repertoire made over the last decade, decoding the individual’s past and present immunological status from TCR repertoire remains a formidable challenge. This is mainly due to the lack of tools to decipher T cell specificity just from the T cell receptor amino acid sequence. This problem involves three components—TCR, peptide, MHC. While the peptide– MHC binding can now be well predicted [36], progress on TCR–antigen binding prediction has been hindered by lack of data, despite some recent progress on the computational side [37–39]. However, direct association between the MHC and the TCR has not been much studied. Our results shed light on this problem by deciphering the HLA restriction of public TCRs found in the repertoires of large cohorts of donors.

An interesting question is whether TCRs with the same HLA restriction share sequence features that would be predictive of MHC binding. Our network analysis carried out through similarity clusters confirms previous intuition [40, 41] that TCRs having similar protein sequences are likely to be specific to the same HLA allele, and we could match some of those clusters to particular viral epitopes. These observations suggest that HLA restriction may be predictable from the TCR sequence alone, which could in turn help refine models of TCR specificity, by accounting for HLA restriction.

Our analysis combines deep bulk *α* and *β* repertoires, allowing us to explore how the alpha chain influences the HLA preference of the TCRs. For some HLA alleles (e.g. B*15:18, DRB3*01:01, DRB4*01:03 or DRB5*01:01), most of the associations we found correspond to alpha clonotypes. However, the *β* chain alone was sufficient to type donors computationally, with little or no improvement upon adding *α* chain information. We expect HLA restriction to be determined by the combination of both chains. Since the repertoire data we used was unpaired, we cannot exclude the possibility that HLA associations learned from paired data could have more predictive power. However, such datasets (deep, chain-paired repertoires of large cohorts of typed individuals) do not exist to the best of our knowledge.

Because the predictive power of our method depends crucially on the number of donors, especially in the HLA-TCR association discovery step, it is limited to common HLA types. Some HLA types are genetically linked, meaning that they tend to occur together. A more general statistical framework could leverage these correlations to improve prediction. Another way to improve statistical power could be to relax the notion of publicness used to allow for imperfect matches (e.g. 1 amino acid different). We showed that this helped the logistic classifier step. Applying this idea to the association discovery step is less straightforward as it would require generalizing the Fisher’s exact test, but it could help increase number of found associations, potentially boosting the predictive power of the classifier.

In addition to our core method, we presented an iterative way to exploit information from untyped repertoires to find new HLA-TCR associations. As the number of publicly available repertoires increases, often from donors for which the HLA type is unreported, this method could prove useful to leverage future datasets to populate expanded lists of HLA-associated TCRs, and to improve the the predictions of computational HLA-typing.

## IV. METHODS

### Datasets

The repertoires initially used for the training of the HLA classifier were extracted from three different datasets described in the main text. The *Russell et al*. dataset [18] is available in The BioProject database under accession code PRJNA762269. The *Emerson et al*. dataset [17] is available at https://clients.adaptivebiotech.com/pub/Emerson-2017-NatGen.

The *Rosati et al*. TCR dataset [19] is available on the ENA database with study accession number PRJEB50045, while HLA genetic data was obtained from the authors upon request. The *Milighetti et al*. validation dataset [21] is available at NCBI Short Read Archive under accession number SUB9362448. All subjects included for the learning phase were typed at 4 digits resolution. For the iterative procedure, the first dataset [34] was downloaded from the NCBI SRA archive using the Bioproject accession PRJNA316572 [34] and the second [35] with donors following a confirmed SARS-CoV-2 infection [35]. From every dataset we excluded donors with incomplete HLA-typing results (i.e. not all loci were known) and duplicated repertoire samples (i.e. coming from different time points) from the same donor Raw sequencing reads were then aligned against a reference database of human V-, D- and J-segments using the MiXCR pipeline [42]. The resulting T cell receptor CDR3 repertoires were further filtered to remove out-of-frame and stop codon-containing CDR3 variants.

### HLA prediction

From the list of HLA-related TCRs, we trained a logistic regressor to predict a given HLA phenotype of an individual from TCR sequencing data.

This input format allows the model to learn and assign different weights to each sequence, maximizing those that are crucial for identifying an allele while minimizing the possible noise introduced by incorrectly identified alleles.

In the logistic model, the probability for a donor to be positive for a particular HLA allele *a* reads:

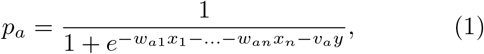

where *x*_1_, …, *x*_*n*_ denote the absence (*x*_*i*_ = 0) or presence (*x*_1_ = 1) of each HLA-associated TCR indexed by *i* in the individual. *w*_*ai*_ are TCR- and HLA-specific weights learned during training. *v*_*a*_ is an extra learnable parameter to encode the information that the alpha chain was sequenced (*y* = 1) or not (*y* = 0) in that individual.

A model was separately trained for each allele *a* on 70% of the typed individuals from the merged cohort (training set) by maximizing the log-likelihood, or equivalently by minimizing the logistic cross-entropy loss, with an L1 penalty which weight was optimized to minimize the loss function on the remaining 30% of individuals (testing set).

## ACKNOWLEDGEMENTS

This work was supported by the Marie Sk-lodowska-Curie Actions H2020-MSCA-ITN-2017 program no 764698 from the European union (AMW, TM, MRO), the European Research Council consolidator grant no 724208 (AMW, TM, MRO), and the Agence Nationale de la Recherche grant no ANR-19-CE45-0018 “RESP-REP” (AMW, TM, MRO), as well as by grants AI136514, AI150747, the St. Jude Center of Excellence for Influenza Research and Response, 75N93021C00016, and ALSAC at St. Jude (PGT). MRO was supported by a grant from the ARC Foundation. PGT has consulted or received travel support from JNJ, Pfizer, Illumina, 10X Genomics, Merck, and PACT Pharma and served on the SAB of Immunoscape, Shennon Bio, and Cytoagents. The funders had no role in study design, data collection and interpretation, or the decision to submit the work for publication.

## SUPPLEMENTARY INFORMATION

**FIG. S1.**
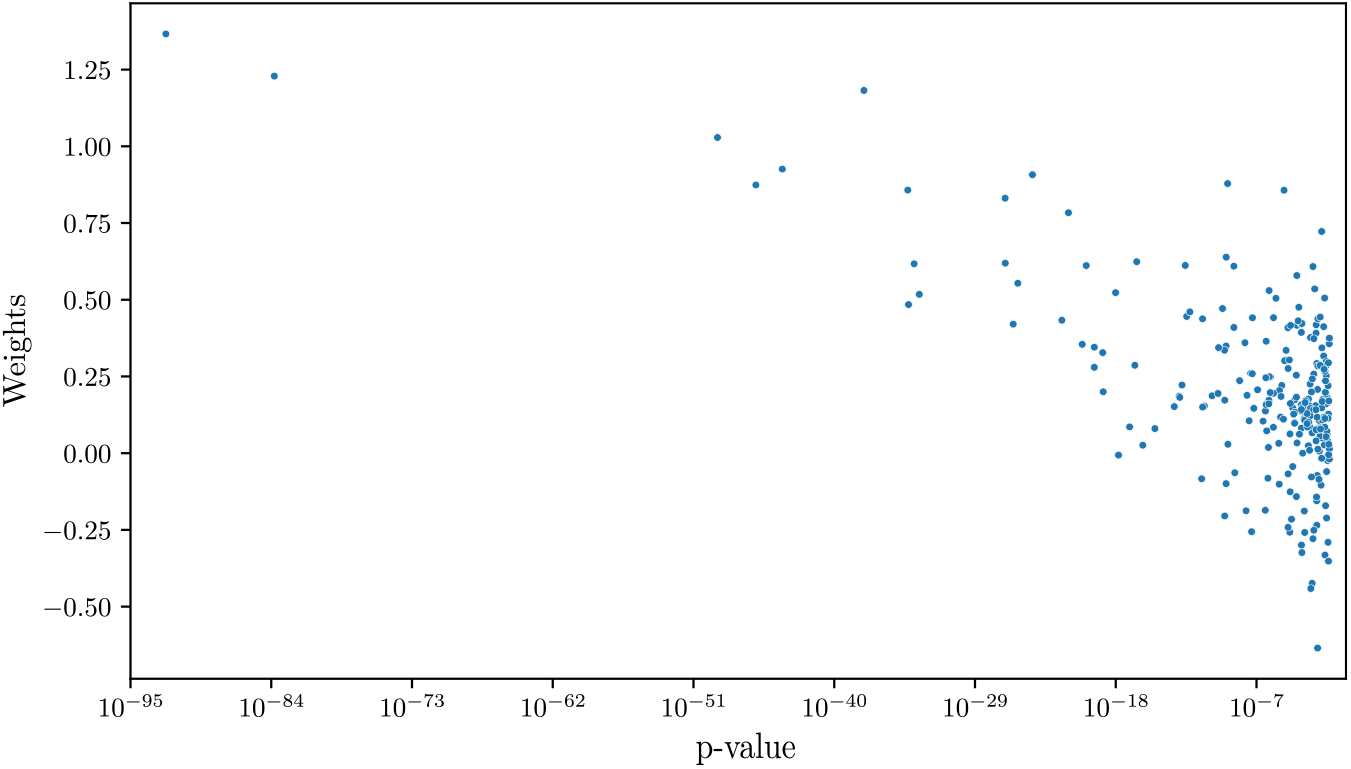
Weights vs. significance comparison. For each of the hits found for the HLA A*02:01, the weight assigned by the logistic regression algorithm (y-axis) is plotted against the significance of the TCR-HLA correlation, i.e. the p-value after Benjamini -Hochberg correction in log scale (x-axis). Pearson coefficient exhibits a remarkable inverse correlation (Pearson coefficient: *−*0.62, p-value:3 *×* 10^*−*28^), confirming the idea that highly HLA-correlated T cell sequences are more helpful to determine the MHC type of the individual.

**FIG. S2.**
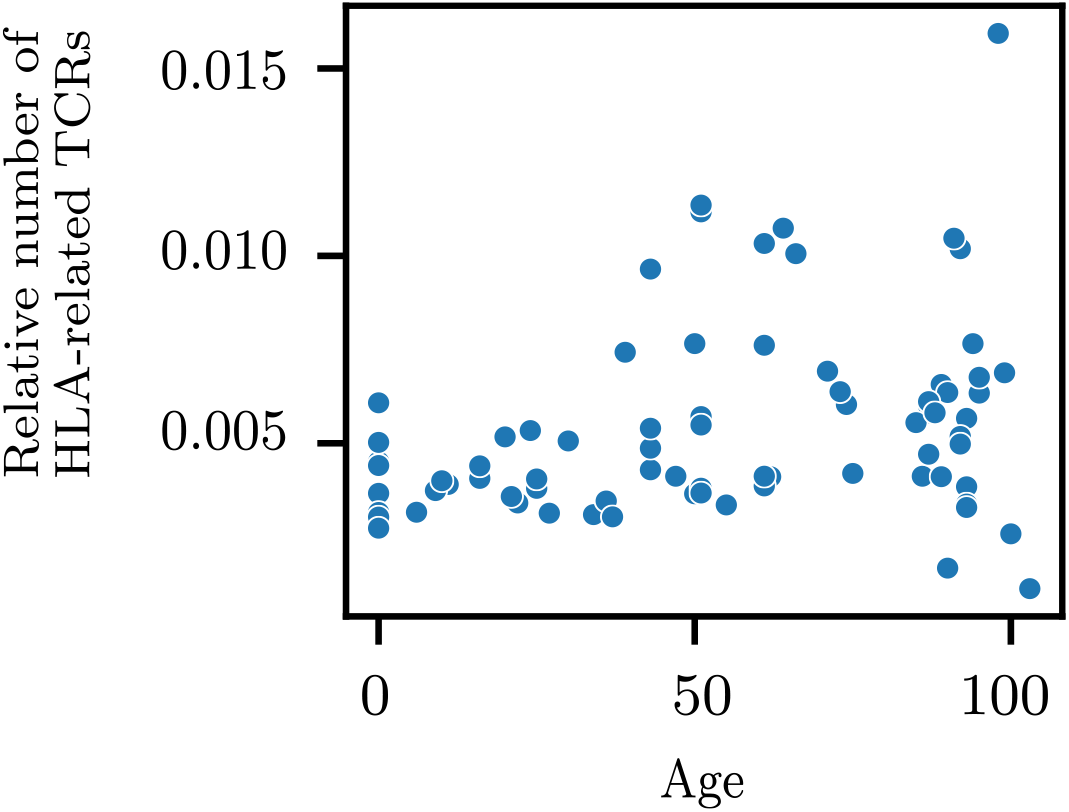
HLA-related TCRs per age range. From *Britanova et al*. [34] the relative number of hits found in every individual is plotted against its age. Despite the suspicion that elder repertoires may cotribute with a higher number of hits due to succesive antigen exposure and selection across lifetime of epitope-responding HLA-restricted TCRs, not a clear correlation is found (Pearson correlation coefficient: 0.29, p-value: 0.01).

TABLE S1. Table of HLA-associated TCRs for both alpha and beta chain, containing their CDR3 amino acid sequence, V gene family, the number of individuals containing the given TCR and the subset of them that also are positive for the corresponding HLA allele, the corrected p-value (*pBH*) that characterizes their association as well as the kind of this last one (if there is a significant overexpression or underexpression of a TCR for the given HLA).

